# Stepwise Evolution and Epistatic Interaction of Driver Mutations from Endometrial Hyperplasia to Carcinoma

**DOI:** 10.64898/2026.06.05.730445

**Authors:** Mary F. Summers, Kira A. Glasmacher, Sem Asmelash, Jeffrey D. Mandell, Nic Fisk, Jeffrey P. Townsend, Vincent L. Cannataro

**Affiliations:** Department of Biology, Emmanuel College, Boston, MA; Program in Computational Biology and Biomedical Informatics, Yale University, New Haven, CT; Yale School of Medicine, New Haven, CT; Department of Cell and Molecular Biology, University of Rhode Island, Kingston, RI, 02881, USA; Program in Genetics, Genomics and Epigenetics, Yale Cancer Center, New Haven, CT; Department of Biostatistics, Yale School of Public Health, New Haven, CT; Department of Ecology and Evolutionary Biology, Yale University, New Haven, CT

## Abstract

To characterize early oncogenesis, pathologically identified pre-cancerous tissue can be analyzed for the presence of cancer drivers. Here, we argue that in such studies, analyses of the driver status of variants, of the association between step-specific prevalence and progression through tumorigenesis, and of driver co-occurrence and mutual exclusivity should be accompanied by estimates of inherent mutation rate of variants and presented within an evolutionary framework of selective epistasis. To illustrate this point, we examine the transition of endometrial tissue from atypical hyperplasia to carcinoma. We apply a step-specific analysis, demonstrating that the strength of selection on somatic driver mutations promoting cell division and survival differs between hyperplasia to carcinoma. We demonstrate that mutations of *PTEN*, which are highly prevalent in carcinomas and have been argued to exert substantial driver effects, exhibit an even larger effect of increasing cellular division and survival within developing hyperplasias. A determination of cooccurrence or mutual exclusivity may be a product of genes sharing or differing in underlying sources of mutation, as opposed to a product of biological interaction and selection. By accounting for tumor-specific mutational processes that influence co-occurrence, we calculate epistatic selective intensities between pairs of drivers. Mutations of *KRAS* and *FGFR2* are often mutually exclusive and were indeed found to exhibit significant antagonistic selective epistasis. However, mutations of *PIK3CA* and *PIK3R1*, which also have been identified as showing mutual exclusivity, do not demonstrate significant antagonistic selective epistasis. Thus, evidence of mutually exclusivity is insufficient to determine epistasis. Accordingly, the application of quantitative approaches that distinctly analyze mutation and selection on cancer variants has the potential to substantially illuminate the trajectory of tumorigenesis and cancer progression.

## Introduction

Understanding which somatic mutations actively promote neoplastic transformation and progression is central to both diagnostic pathology and precision oncology. Variant prevalence has often been used as a metric of their biological and clinical importance [1]. However, prevalence reflects both selective advantage conferred and also the intrinsic rate at which variants arise *in vivo* in individual cancer-competent cell lineages, i.e. the cellular mutation rate [2], which can vary by more than an order of magnitude among genes [3]. Evolutionary analysis addresses this conundrum by comparing observed substitution rates with neutral expectations, thereby estimating the strength of selection conferred by specific variants [4]. These estimates clarify which mutations promote clonal proliferation or survival and can inform interpretation of therapeutically targetable variants [5]. With expanding sequencing datasets, including data from precursor and malignant lesions, this framework can now be applied to define how somatic mutation rates, selection intensities, and driver-gene interactions change across histopathological progression.

In endometrial tumorigenesis, atypical hyperplasia (AH; also termed endometrial intraepithelial neoplasia [6]) is widely regarded as a direct precursor to endometrial carcinoma (EC). Approximately one third of patients diagnosed with AH subsequently develop EC [7], and recent research has demonstrated shared mutations in adjacent AH and EC lesions, supporting clonal continuity [8].

Li *et al*. further compared the five most commonly mutated genes in AH and EC, identifying *PTEN, PIK3CA/PIK3R1*, and *ARID1A* as recurrent drivers enriched in carcinoma relative to paired precursor lesions [8]. They proposed that accumulation of these mutations may promote progression from AH to EC. Subsequent studies have likewise used variant prevalence, co-occurrence, and mutual exclusivity to infer differences in somatic drivers of AH and EC [9,10]. However, these analyses did not account for underlying mutation rates or quantify stepwise selection during endometrial tumorigenesis. Here, we estimate the strength of selection—the relative cell-proliferative or cell-survival benefit conferred by a mutation, or the cancer effect size—to identify genetic changes most likely to drive progression from hyperplasia to carcinoma.

## Methods

### Data sources

We analyzed published whole-exome sequencing data from AH (*n* = 30) and Stage I EC (*n* = 513). AH data were obtained from Li et al. [8], Stage I EC data were obtained from Dou et al. [11] as well as The Cancer Genome Atlas (TCGA), accessed through the Genomic Data Commons Data Release 45 [12,13].

### Driver-event definitions

Driver mutations were defined in eight genes previously implicated in endometrial carcinoma: *PTEN, PIK3CA, PIK3R1, ARID1A, CTCF, CHD4, KRAS*, and *CTNNB1 [8]*. We also included driver mutations identified in *FGFR2*, a potential epistatic partner of KRAS [14]. Genes were classified as oncogenes or tumor suppressor genes according to OncoKB [15]. Recurrent variants were considered as possible driver events in the oncogenes *PIK3CA, CHD4, KRAS, FGFR* and *CTNNB1*. Recurrent variants, nonsense variants and truncating variants were all included in evaluations of the tumor suppressor genes *PTEN, PIK3R1, ARID1A*, and *CTCF*. Variants meeting these criteria were aggregated by gene for selection inference.

### Step-specific mutation rates and selection inference

We used a framework in cancereffectsizeR (v2.10.2, [16]) to estimate neutral mutation rates and quantify selection on somatic driver events across two steps of progression. This framework compares observed driver-event frequencies with expectations under neutral somatic evolution, separating the probability that a variant arises within a cell from the selective advantage conferred to that cell within a tissue. Selection was estimated separately for Step 1, progression from normal endometrial precursor cell lineages to AH, and Step 2, progression from AH to Stage I EC. This design followed a step-specific framework previously applied to esophageal tumorigenesis [17].

For selection inference, each sequenced sample was treated as representing a dominant clonal expansion. Although individual biopsies may contain subclonal heterogeneity, variants detected by bulk whole-exome sequencing are expected to be present at sufficient cellular frequency to reflect expansion within the sampled lesion. Accordingly, observed variants were modeled as substitutions that had arisen and become established within the sampled neoplastic or pre-neoplastic cell population.

To test whether selection differed significantly between steps, we compared nested models for each gene: a full step-specific model that provided distinct estimates of selection on variants along the normal-to-AH and AH-to-EC transitions, versus a reduced model providing a single estimate of selection parameterizing to both transitions. Statistical significance was assessed using likelihood-ratio tests against a χ2 distribution with one degree of freedom.

### Pairwise selective epistasis

To evaluate whether the selective effect of one driver depended on another, selection on each driver in a pair was estimated in the absence or presence of its partner [16,18]. Reduced selection in the mutant background was interpreted as antagonistic epistasis; increased selection was interpreted as synergistic epistasis. Evidence for epistasis was assessed by likelihood-ratio tests comparing a model that allowed selection to differ by mutational background with a nested model without mutation-dependent selection.

Because co-mutations of driver genes are relatively rare, and because epistasis often manifests across time as mutations arise, the pairwise epistasis analysis was performed across the full available sequenced cohort rather than separately within each progression step, maximizing power to detect mutant-background-dependent changes in selection.

## Results and Discussion

### An evolutionary framework enables quantitative assessment of how driver mutations contribute to stepwise progression in endometrial tumorigenesis

Several alterations, including those in *PIK3CA, ARID1A*, and *PIK3R1*, and *FGFR2*, were more frequent in Stage I EC than in AH (**Fig. 1**). This pattern suggests continued selection during progression to carcinoma. However, prevalence alone cannot determine which mutations are favored or how much, because mutation rates and mutational processes may differ between steps. Distinguishing altered selective advantage is therefore essential for interpreting driver enrichment during progression from AH to EC.

**Figure 1:**
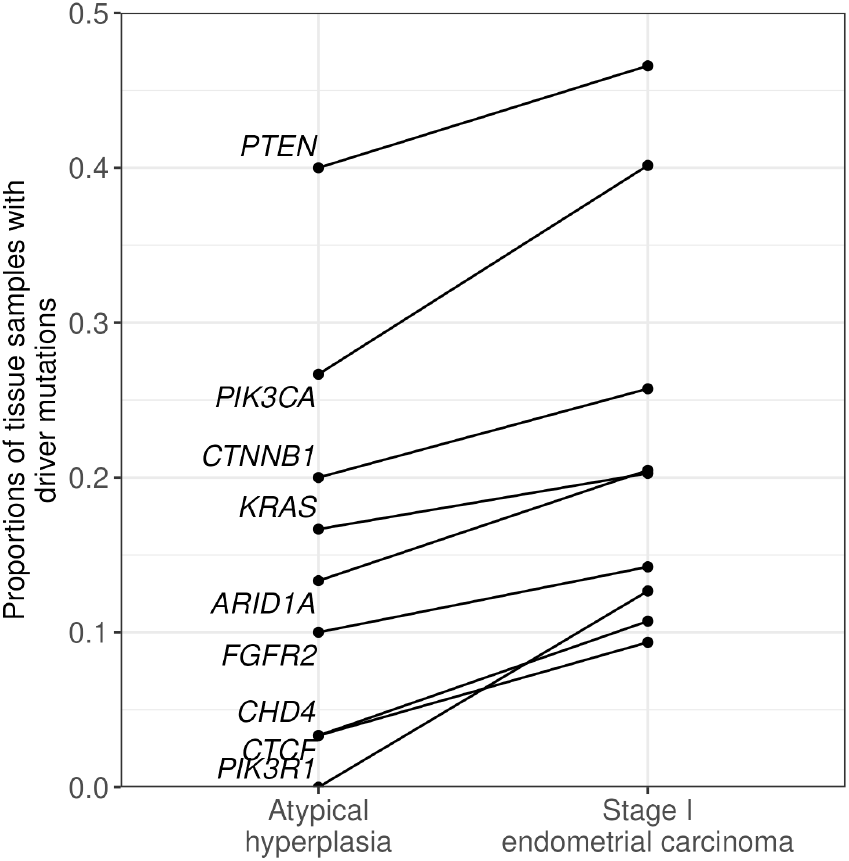
Proportions of tissue samples harboring driver mutations in eight genes reported in Li et al. [8] as well as *FGFR2* in atypical hyperplasia (n = 30) and Stage I endometrial carcinoma (*n* = 513).

Analysis of nine genes revealed differential selection between the normal-to-AH and AH-to-Stage I EC transitions (**Fig. 2**). *PIK3R1, CTCF, CHD4, ARID1A, PIK3CA*, and *FGFR2* experienced stronger selection during progression from AH to Stage I EC, indicating that selection continues beyond the hyperplastic stage. Notably, PIK3CA mutations were more prevalent in carcinoma than AH (**Fig. 1**), but did not show a significant increase in selection strength, suggesting that increased mutation rates rather than oncogenic selection explain the increased frequency of *PIK3CA* variants in EC. In contrast, *CTNNB1, PTEN*, and *KRAS* showed strong positive selection during progression from normal tissue to AH (**Fig. 2**), consistent with early driver selection in cancer [19,20]. Likelihood-ratio tests comparing a model estimating distinct selection coefficients for the two steps of progression to a model with a single shared coefficient confirm that the two-step model is a significantly better model of evolution for *PIK3R1*, pointing to both the strong increase of selection for mutations within this gene after tumorigenesis and also the need for additional sequencing data, especially from premalignant tissue, to resolve additional relationships.

**Figure 2:**
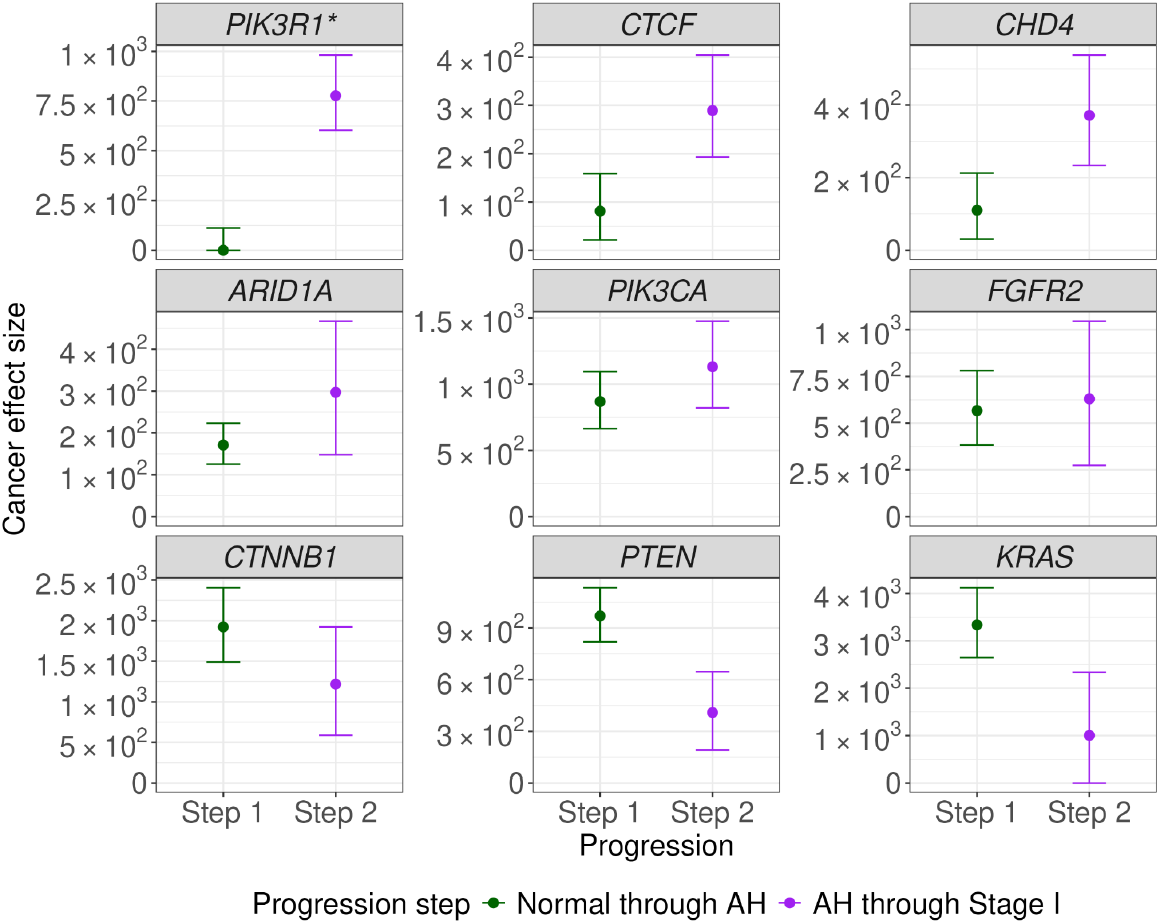
Cancer effect size (strength of selection; increases in cell division and/or survival) and 95% confidence intervals (error bars) on nine genes in precursor cells to hyperplasia (Step 1, filled green circles) and in cell lineages evolving from hyperplasia (Step 2, filled purple circles) into Stage I endometrial carcinoma. Statistical significance of step-specific differences in selection are based on likelihood-ratio tests comparing a model with separate selection coefficients for the two steps of progression to a model with a single shared coefficient (^*^: *P* ≤ 0.05).

### Observations of mutational exclusivity among variants should be evaluated with quantifications of epistasis

Changes in selection during tumor progression likely reflect in part the evolving genetic context of the neoplasm, including interactions among driver mutations [18]. Such interactions are often inferred from mutual exclusivity, which is commonly interpreted as evidence of functional redundancy or antagonistic pathway effects. Li et al. [8] and Levine et al. [12] reported mutual exclusivity between mutations of the known interactors *PIK3CA* and *PIK3R1*, and Russo et al. [14] reported similar patterns for *KRAS* and *FGFR2*. Yet TCGA includes tumors with co-mutations in these same gene pairs, indicating that exclusivity is not absolute. These rare but present co-mutations clarify that it is important to analyze datasets with substantial sample sizes, quantifying underlying mutation rates when evaluating genetic dependencies.

To assess how genetic interactions influence selective pressures during tumor progression, we quantified selection for *PIK3CA*–*PIK3R1* and *KRAS*–*FGFR* driver events in the absence and presence of mutation in each partner gene. The analysis revealed a trend of selective epistasis between mutations of regulatory subunit–catalytic partner genes *PIK3CA* and *PIK3R1*, though not statistically significant (**Fig. 3A**). In contrast, selection for *FGFR2* mutation was statistically significantly reduced in the presence of downstream *KRAS* mutation, reaching levels indistinguishable from neutrality (**Fig. 3B**). This substantial attenuation is consistent with antagonistic epistasis, in which alteration of one driver reduces the selective benefit of altering the other. The result accords with canonical RAS-MAPK pathway architecture, where *FGFR2* and *KRAS* act within the same signaling cascade and may therefore be functionally redundant when both are mutated [21]. Thus, mutual exclusivity should be treated as an indirect indicator rather than as *prima facie* evidence of epistasis. Drawing conclusions from mutual exclusivity should be avoided; instead, quantitative estimates of selection should instead be interpreted in the context of intrinsic mutation rate, progression step, and driver-gene context.

**Figure 3:**
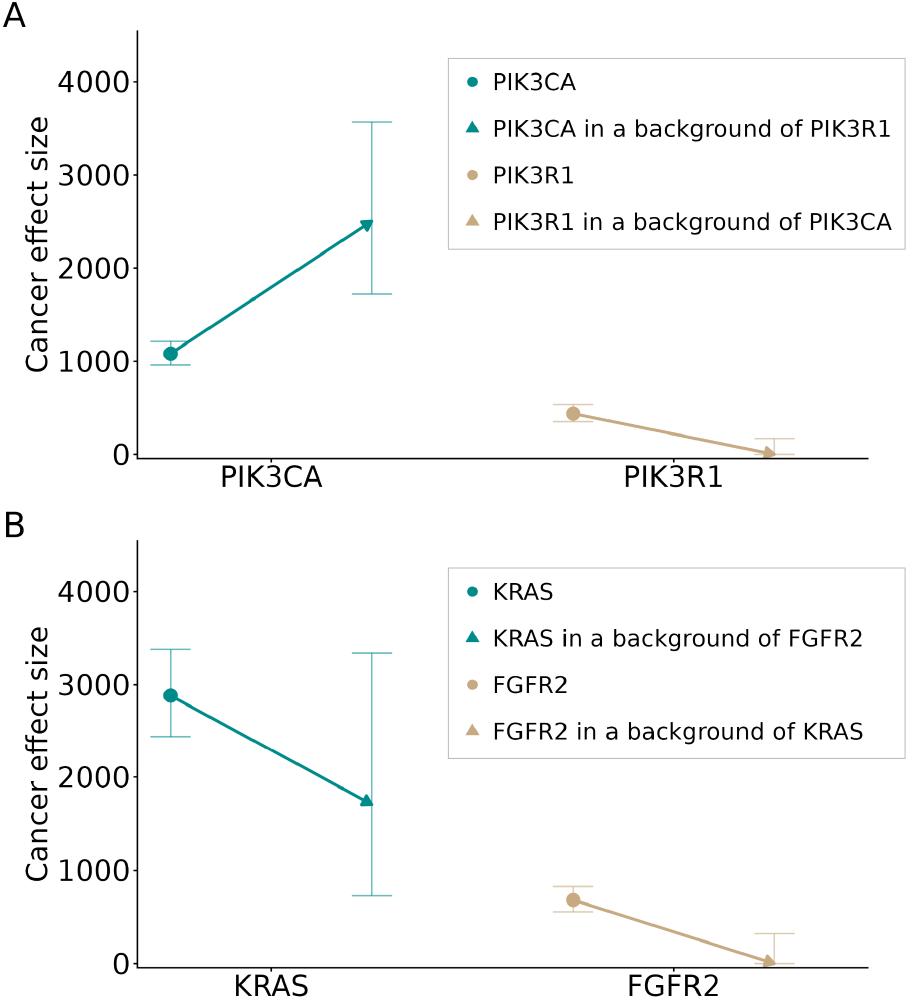
Quantification of cancer effect (strength of selection) and pairwise epistatic effects. (**A**) Selection on mutations in *PIK3CA* (teal point) appears increased in a background of PIK3R1 (teal arrowhead), and selection on *PIK3R1* (gold point) is decreased in a background of mutated *PIK3CA* (gold arrowhead; these epistatic interactions are not statistically significant: *P* = 0.13, likelihood ratio test), and (**B**) Selection on mutations of *KRAS* (teal point) is decreased in a background of mutated *FGFR2* (teal arrowhead), and selection on *FGFR2* mutations (gold point) is decreased in a background of mutated *KRAS* (gold arrowhead; these epistatic interactions are statistically significant: *P* < 0.001, likelihood ratio test).

Because mutation and subsequent clonal expansion within a neoplastic lineage are uncommon, co-mutation of major drivers is observed infrequently. Reliable detection of epistatic interactions therefore requires large tumor sequence data sets, and even strong epistatic interactions may be difficult to quantify when driver mutation rates are low. These challenges justify caution when interpreting mutual exclusivity among cancer drivers in small cohorts, where statistical signals may reflect correlated mutational processes rather than selective interaction or known biological function.

### Sequencing of histologically normal tissue is essential for fine-scale reconstruction of the process of tumorigenesis

Normal and premalignant sequence data remains limited compared to replete tumor sequence datasets for deeply researched cancer types. Future studies should make use of opportunities to sequence histologically normal and premalignant endometrial tissue. Expanding such reference data sets will enable more accurate comparison of mutation rates and selective pressures between normal, hyperplastic, and malignant states. Additional data will enable identification of early driver mutations, clarification of the steps at which they exert selective advantage, and revelation of the mutational backgrounds that promote progression to carcinoma.

## Reproducibility

Code to reproduce the analyses conducted within this manuscript is found at https://github.com/Cannataro-Lab/endo_stage_epi_paper.

